# *CaneCestry:* A Web-based Toolbox for Efficient Pedigree Analysis and Visualization

**DOI:** 10.1101/2025.05.02.651868

**Authors:** Zachary Taylor, Brayden Blanchard, Ajay Dhungana, Collins Kimbeng

## Abstract

Genetic variability is the lifeblood of all breeding programs. This variability is acted on via selection to identify elite progeny that can be utilized agronomically or exploited as parents. Sugarcane breeders are tasked with creating optimal genetic variation through crossing. Prior knowledge of pedigree and parental performance allows breeders to discern which crosses to prioritize, since the time to make crossing decisions and the space to evaluate tens and thousands of progenies are both limited resources in sugarcane variety development programs. In this project, we have developed a user-friendly, web-based platform called *CaneCestry* that provides a wide range of tools utilizing pedigree information. *CaneCestry* can be utilized to generate family trees for parents involved in a potential cross, enabling the breeder to efficiently display and visualize the lineage of the genotypes contained within. Kinship matrices are a powerful numerical representation of relationships between individuals in a population. While the calculation of kinship can be computationally intensive, *CaneCestry* is built upon a framework allowing for efficient cloud-based computation. These numerical representations of kinship elucidate the relatedness of potential parents, allowing the breeder to make the most informed crosses while avoiding inbreeding depression. Kinship matrices can also be utilized to enhance prediction models by adding the relatedness of the involved genotypes as a covariate, accounting for additive genetic control of traits. *CaneCestry* provides a comprehensive, web-based set of tools available in the office and field, enabling sugarcane breeders to utilize pedigree information in all stages of their breeding program.

## Introduction

Genetic variability is the lifeblood of all variety development programs. Primarily, but not exclusively, created through crossing; this variability is acted on via selection to identify elite progeny that can be utilized agronomically or exploited as parental lines [1]. This process leads to a cyclical pattern of making crosses, evaluating progeny, and taking noteworthy members of those progeny populations and crossing them. Over time, these processes precipitate an abundance of information on the crosses that have been made and their resulting progeny. This pedigree data can then be exploited to the benefit of the breeder, enhancing decision-making by providing information on relationships within and between populations.

Two key tools for analyzing pedigree data is the numerator relationship matrix (A Matrix), which calculates the additive genetic relationship between individuals [2], and the Kinship matrix, being the probability that a given allele is identical by descent between individuals[3]. These probabilities can be used as a numerical representation of the relatedness between genotypes. This information, in numerical format, can enhance statistical modeling approaches with the relationship between individuals in breeding trials and other studies considered as their covariances [4]. Relatedness of individuals can also be utilized to greatly enhance crossing decisions, enabling breeders to control or exploit inbreeding in their populations[5].

Several software options, such as Helium [6] and R packages like AGH matrix [7], currently exist to visualize pedigrees and calculate relationship matrices, respectively. However, they are inadequate in their handling of large and complex historical pedigree datasets. Historical pedigree datasets spanning decades to centuries can be extremely large, leading to increased computational time and potential errors [8] while calculating these matrices. Current tools act on an assumption of data pre-processing (subsetting and cleaning) to combat these limitations. These methods act on entire datasets without allowing the user to specify individuals of interest, which can exponentially increase runtime and memory usage when calculating matrices from large historical datasets.

Graph theory is the subsect of discrete mathematics focused on graphs, or pairwise relationships between objects modeled as nodes and edges. Due to the tree-like structure of pedigree data with nodes representing individuals and edges representing relationships, directional acyclic graphs (DAG) [6] have been historically used to mathematically interpret these data. One of the most common algorithms to search or quantify relationship distance between nodes is breadth-first-search (BFS) [9]. Breadth-first-search is an efficient method to explore pedigree data as it starts from a specific individual (root node) and visits all immediate relatives (connected by one edge, children or parents) before moving to the next generation. Generations are iteratively visited until no unexplored nodes remain, at which point the search ends [9]. Thus, this study overcame pre-processing challenges by integrating a graph traversal–based subsetting method, employing BFS to incrementally traverse the complex family network of selected individuals. This targeted traversal omits any individuals not relevant to the individual(s) of interest, circumventing high computation load or data pre-subsetting.

To provide a more user-friendly approach to pedigree analysis, we developed *CaneCestry*, an all-in-one web-based platform for pedigree dataset visualization and manipulation for complex historical sugarcane pedigrees. Through an integrated approach that incorporates our proposed graph traversal–based subsetting method into its core functionality, *CaneCestry* rapidly generates numerator relationship matrices and family trees using cloud-based computation. This provides faster and reliable access to critical pedigree insights regardless of coding expertise or available computational power, hence enabling breeders to exploit comprehensive information to inform their breeding and selection decisions.

## Design and Implementation

*Canecestry* is a web-based toolbox that allows plant breeders to utilize large pedigree datasets conveniently and efficiently. This is done through a web-based approach, ensuring that all computationally challenging tasks are completed in the cloud. Computational complexity is further mitigated using our proposed graph traversal subsetting technique, ensuring that only individuals necessary to the interest of the breeder are utilized. This addresses issues present in existing tools, as most rely on datasets that have been pre-processed. *Canecestry* is currently hosted through pythonanywhere.com at (https://canecestry-zanthoxylum2117.pythonanywhere.com/ [Accessed 24 Feb 2025]). This software has been designed to work with popular browsers (Chrome, Edge, Safari), as well as on mobile devices. *CaneCestry* can also be run locally using Docker, or the environment available for download on the GitHub page. Standard format for a pedigree dataset is used, with Line, Male Parent (sire) and Female Parent (dam) being the column names. Data in the form of tab delineated text (.txt) file is currently the singular format supported.

*Canecestry* is built with Python 3.9, utilizing the Dash [10] and Flask [11] web frameworks. It leverages Pandas 1.3.5 for data manipulation and storage, while NumPy 1.21.6 [12] handles numerical computations. Graphviz 0.20 [13] is used to generate pedigree graph visualizations, and Seaborn 0.11.2 [14] is employed for creating heatmaps.

Features are presented in a modular format, with the user first being prompted to upload their own data or use the pre-loaded sugarcane pedigree dataset. Thereafter, users are able to select a method to calculate numerator relationship matrices, explore the pedigree data, or append new entries. The functionality to append new entries into the dataset is primarily useful for sugarcane breeders, as the preloaded sugarcane dataset is comprehensive and housed in the cloud. Functionalities and features will be explained and showcased in the following sections.

### Sugarcane Data

Canecestry was developed and subsequently tested using a historical U.S. sugarcane pedigree dataset containing 49,163 individuals. Due to the nature of how sugarcane is bred, its pedigrees exhibit great complexity and redundancy [15]. Modern sugarcane cultivars are complex aneu-polyploid hybrids resulting from decades of introgression from wild relatives and recurrent selection. Cytotypic variation is extensive between cultivars, with each homologous group (HG) (ie chr1) potentially containing variable chromosome copies and not holding to a consistent ploidy across all HG’s. [16]. These HG’S contain varying numbers of homeologous chromosomes, with these groups being comprised of S. *officinarum*, S. *spontaneum*, interspecific recombinant, or any combination of these chromosome origins[17]. These progenitor species are highly polyploid containing both S. *officinarum* (2n = 8x = 80, x=10) colloquially known as the “Noble” cane, and S. *spontaneum*, displaying extensive cytotypic variation [18] (tetraploid to dodecaploid) with a base chromosome count range of x = 8,9,10 due to chromosome fusion [19] or descending dysploidy [20]. Additional contributors to the gene pool include cultivated “species” (hybrids) [21] such as S. *barberi* (originating in India) and S. *sinense* (originating in China), some of the earliest documented hybrid cultivars between S. *officinarum* and S. *spontaneum* [22]. The putative progenitor of S. *officinarum*, S. *robustum* (x = 10, 2n = 6 − 20x = 60 − 200) [23], provides valuable diversity [22] and disease resistance, while contributing to the cytogenetic complexity. Due to this complexity, genomic information is not as readily available for sugarcane breeders, with currently only two genome sequences being published to date[16, 17].

Sugarcane is primarily bred via biparental crossing, due to the high amount of heterozygosity and cytotypic variation between hybrids[24]. Most sugarcane hybrids descend from a pool of around 10 progenitor genotypes [25] from the noble cane S. *officinarum*, or wild relatives S. *spontaneum* and S. *robustum*. Early interspecific crosses between S.o and S.s exhibited second division restitution [26], or 2n+n inheritance, leading to this early genomic complexity. Historical crosses leading to this diverse genetic background perpetuated this complexity, with abnormal patterns of inheritance and aneuploidy leading to drastically different numbers of homeologous groups (subgenomes) within each HG of each sugarcane hybrid[17]. Due to resource limitations, biparental crosses are not always feasible, leading to the practice of using multiple pollen parents (males) for a single female parent [27]. These ambiguous crosses, known as polycrosses, obfuscate the paternity of the progeny from such crosses without the use of DNA fingerprinting [27], which can be resource-draining and time-consuming. Uncertainty of paternity from polycrosses can hinder the usage of these pedigree-based tools by breeders, as historically they have been listed as a male parent under an arbitrary identifier. *Canecestry* is built on a framework that mitigates these pedigree dead ends by treating a polycross as a unique dead end to avoid inflating the relatedness of progeny resulting from a single polycross. This is accomplished by utilizing the consistent nomenclature used by sugarcane breeders to denote polycross parentage and omitting them from calculation.

The dataset used in this study contains 2,631 unique female parents and 2,132 unique male parents of commercial hybrid or wild accession origin.

### Data Subsetting

*Canecestry* was designed for efficient, user-friendly manipulation of large historical pedigree datasets. While tools currently exist to extract family trees and additive kinship matrices from these datasets, preprocessing is required for effective usage when large historical datasets are used. Canecestry employs a graph traversal-based subsetting method, utilizing breadth-first search to include only individuals relevant to the user’s interest. As outlined in **Figure 1**, the workflow of this subsetting algorithm for the additive kinship matrix takes an individual or list of individuals provided by the user. Lists of individuals can be input using the dropdown menu or as a comma-delineated string. A secondary list (visited set) is generated at this time for individuals that have been “visited”, and the individuals present in the initial queue are added to the list. The members of the queue are then used to search the full dataset to identify parents and children, which are subsequently added to the queue and, once visited, to the visited set (list). This process continues until no further parents or children can be identified. During this process, a topological sorting function is also employed to ensure the placement of individuals in the order of mother-father-child. This sorting process ensures the correct calculation of additive kinship matrices. The Visited list, which contains the completed subset of the full dataset, is used to perform the chosen analysis. This workflow is the same as the subsetting algorithm used for family tree generation, except that descendants are neither searched nor added to the visited list. This ensures that the base of the pedigree visualization consists only of the individual of interest.

**Figure 1.**
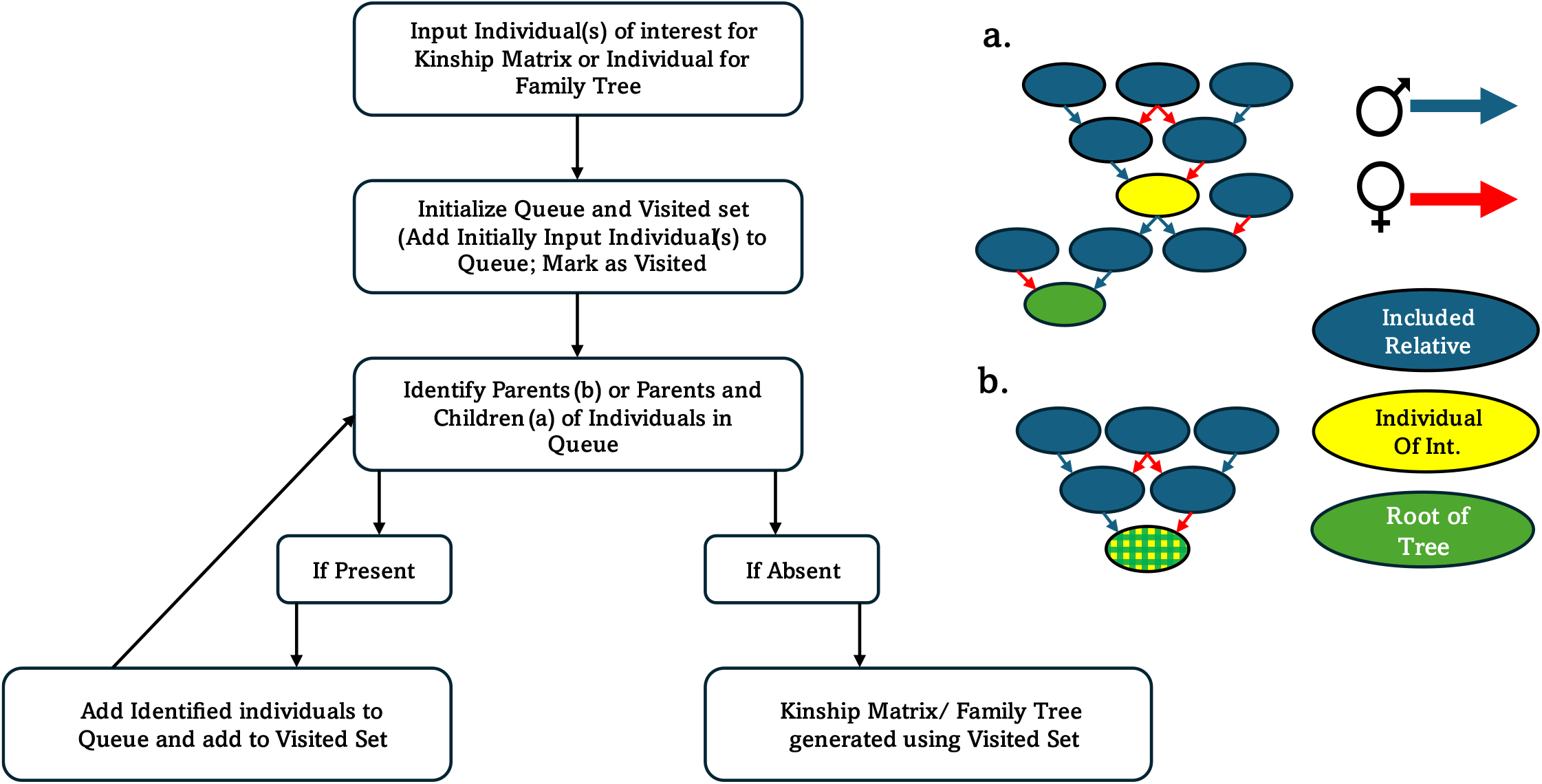
Workflow of pedigree subsetting algorithm. Canecestry employs a breadth-first search algorithm to subset the input pedigree dataset to only include individual(s) of interest and their relatives. Two methods of defining relevant nodes are employed with (a) including ancestors and progeny, and (b) only including ancestors.

### Matrix Calculation

Additive kinship matrices are calculated using the recursive method outlined by Henderson[2]. Each pedigree entry in the now subset pedigree data is iterated over. If no parents are known, the individual is assigned a diagonal value of one. If one parent is known, the diagonal value is set to one and the off-diagonal elements are half of the known parent’s relationship values. When both parents are known the diagonal entry for the individual is 1 + 0.5 × *A*(*mother, father*) and each off-diagonal element is half the sum of the mother’s and father’s values. For the off-diagonal elements, the computation is given by:

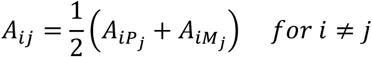

where 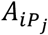 is the pedigree relationship between individual i and the female parent of individual j, and 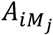 is the pedigree relationship between individual i and the male parent of individual j. Each row in the pedigree is processed iteratively to update the matrix. The matrix is stored as a NumPy array and converted to a labeled data frame for indexing.

The full matrix is available to the user to download as a .csv file format. The user can also visualize a heatmap and dendrogram of the matrix. This visual can be used to identify if there were individuals listed that had separate lineages with no relation to one another or groups of highly related individuals. The user is presented with the option to subset the matrix as well, choosing from a dropdown list of individuals to include. This feature is an optimization for users utilizing additive kinship matrices in a setting where only certain values are necessary, as these matrices from historical datasets can be hundreds of megabytes to a gigabyte or more in size.

As outlined in **Figure 1**, two methods are presented to the user to calculate these matrices. The first method (a) considers both the ancestors of the given individual(s) as well as the progeny. This precipitates a more comprehensive matrix of the population relating to the individuals and can be useful to quantify the genetic contribution of a genotype; however, it can be computationally intense if many of the listed individuals have many offspring. The second method (b) only considers the ancestors, leaving the selected individuals as the “root” of the tree. This allows for faster computation when the user only needs the relationships between the given individuals of interest. Shown in **Figure 2** and **Figure 3** are heatmaps of an example of each method. Method (a) took 43.52 seconds to run and compile all ancestors and progenies of the two individuals, while method (b) took 6.23 seconds only. These matrices were generated using two of the sample genotypes in the sugarcane dataset L01-0299 and HoCP14-0885. A snippet of the numerical output matrix is shown in **Table 1**, demonstrating the output of this information with the sample genotypes and their parents.

**Table 1.** Subset Numerator Relationship Matrix (A Matrix) of L01-0299 and HoCP14-0885 and their Parents. Numerator relationship matrices can be subset, to provide ease of use for users, by cutting down on the size of the output matrix while retaining all information. Highlighted values are the relationships of the individuals with themselves, or 1+ the coancestry of the parents.

|  |  |  | Male Parent 885 | Female Parent 885 | Male Parent 299 | Female Parent 299 |
| --- | --- | --- | --- | --- | --- | --- |
|  | **L01-0299** | **HoCP14-0885** | **Ho07-0613** | **HoCP05-0920** | **LCP85-0384** | **L93-0365** |
| **L01-0299** | 1.153382 | 0.310191 | 0.430518 | 0.189864 | 0.734638 | 0.720093 |
| **HoCP14-0885** | 0.310191 | 1.097925 | 0.680914 | 0.60993 | 0.383873 | 0.236509 |
| **Ho07-0613** | 0.430518 | 0.680914 | 1.165978 | 0.19585 | 0.580453 | 0.280583 |
| **HoCP05-0920** | 0.189864 | 0.60993 | 0.19585 | 1.02401 | 0.187293 | 0.192435 |
| **LCP85-0384** | 0.734638 | 0.383873 | 0.580453 | 0.187293 | 1.162512 | 0.306765 |
| **L93-0365** | 0.720093 | 0.236509 | 0.280583 | 0.192435 | 0.306765 | 1.133422 |

**Figure 2.**
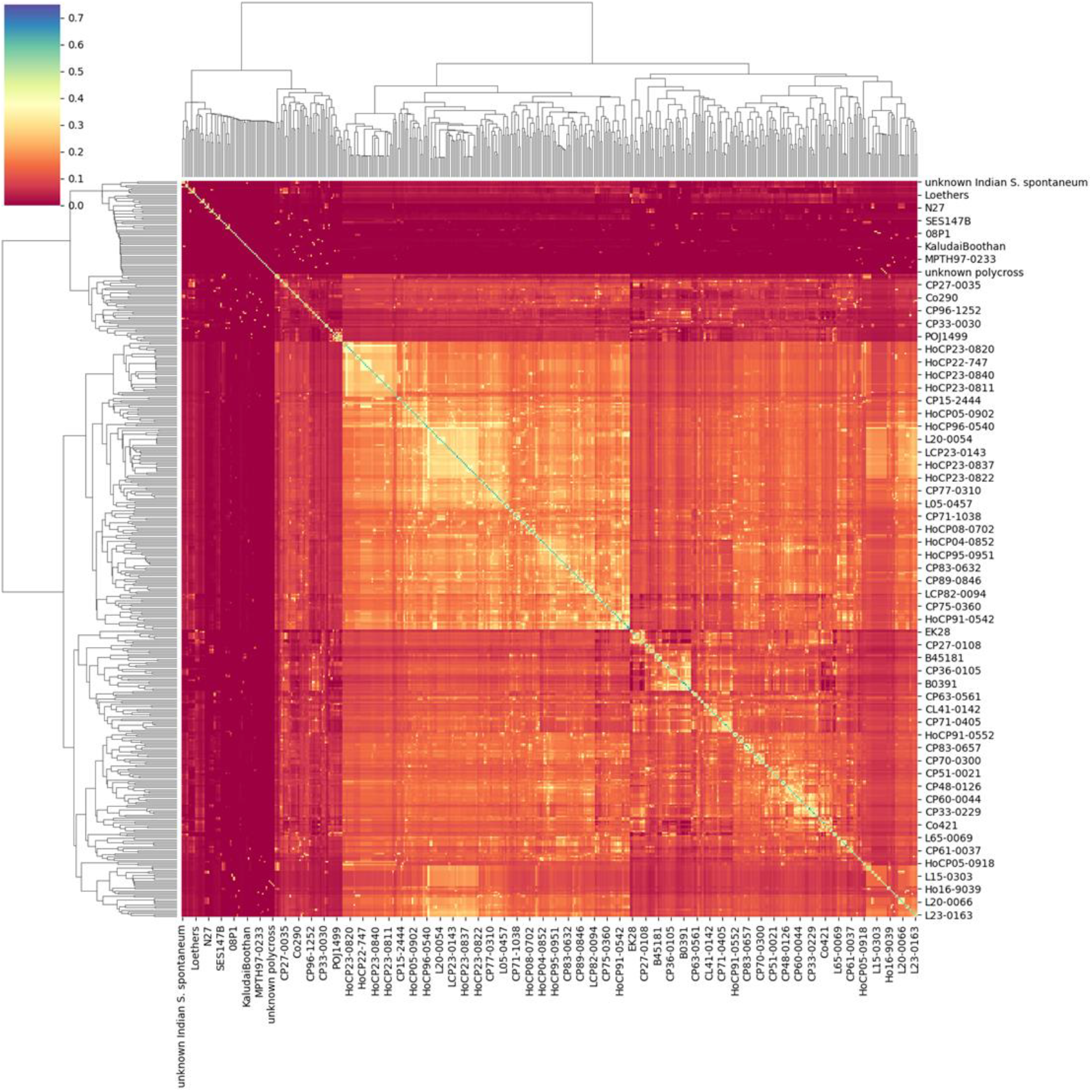
Heatmap from numerator relationship matrix of sugarcane genotypes L01-0299 and HoCP-0885, using subsetting method a that contains all ancestors and progeny of the two genotypes of interest. Axis Variety names are only a selected few of the total number of individuals and are not representative of the total number included in analysis.

**Figure 3.**
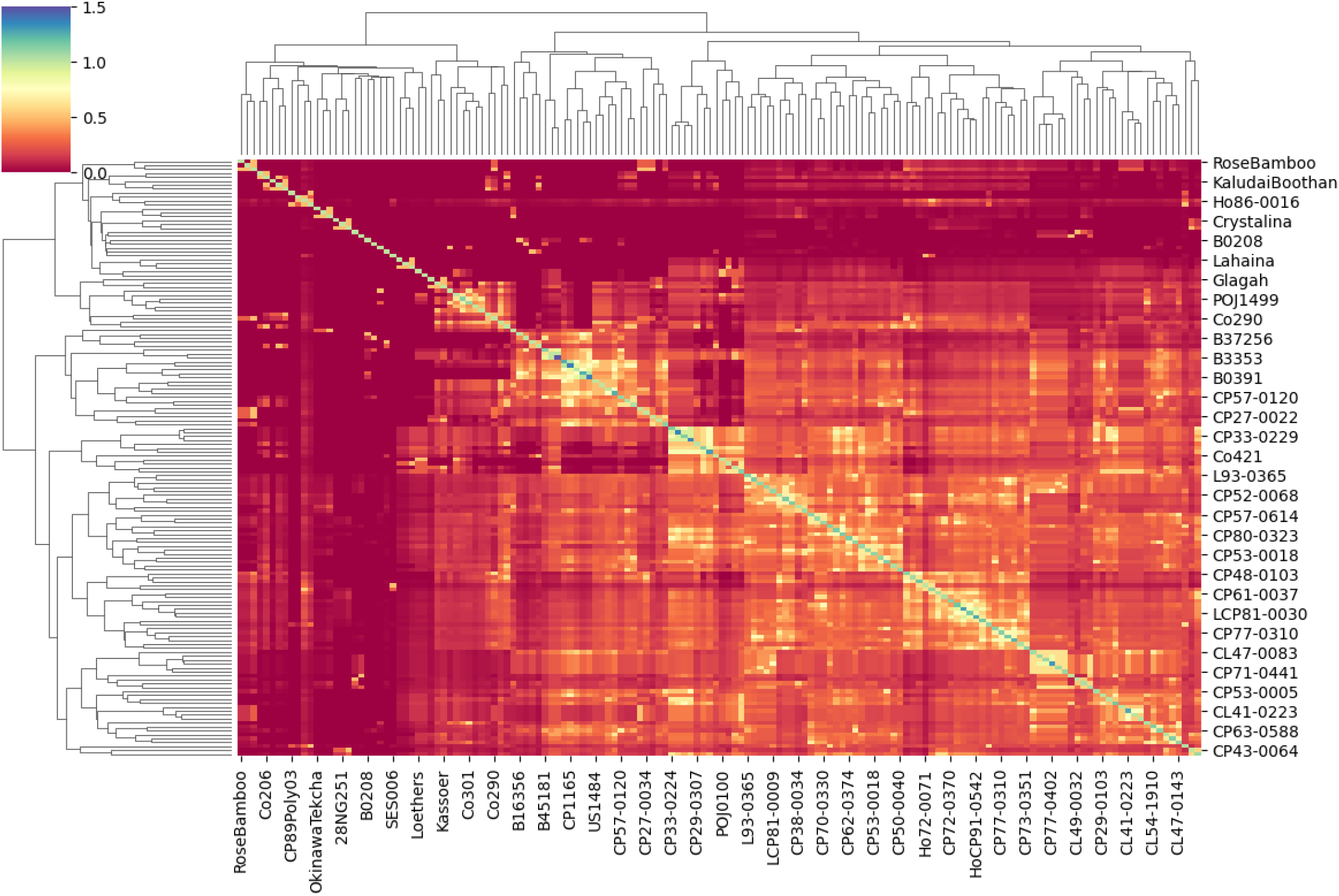
Heatmap from numerator relationship matrix of sugarcane genotypes L01-0299 and HoCP-0885, using subsetting method b that results in a matrix of just the ancestors of the two individuals of interest. Axis Variety names are only a selected few of the total number of individuals and are not representative of the total number included in analysis.

### Family Tree Generation

Pedigree visualizations are a core functionality of *Canecestry*, providing a convenient display of the ancestry of genotypes of interest. Pedigrees are displayed as DAGs or directed acyclic graphs, as previously utilized in other pedigree software such as Helium. An additive kinship matrix is computed at the same time as the family tree is generated, and the relatedness between an ancestor and the specified genotype is added to the individual’s node on the tree. Nodes are generated with a color scheme to highlight the relatedness between the specified individual and each node and are accomplished using the calculated additive kinship values. This feature is present to ensure confidence in calculated values, as well as a tool to understand these values. An example pedigree of L01-0299, a highly utilized cultivar in the Louisiana sugarcane industry, is presented in **Figure 4**.

**Figure 4.**
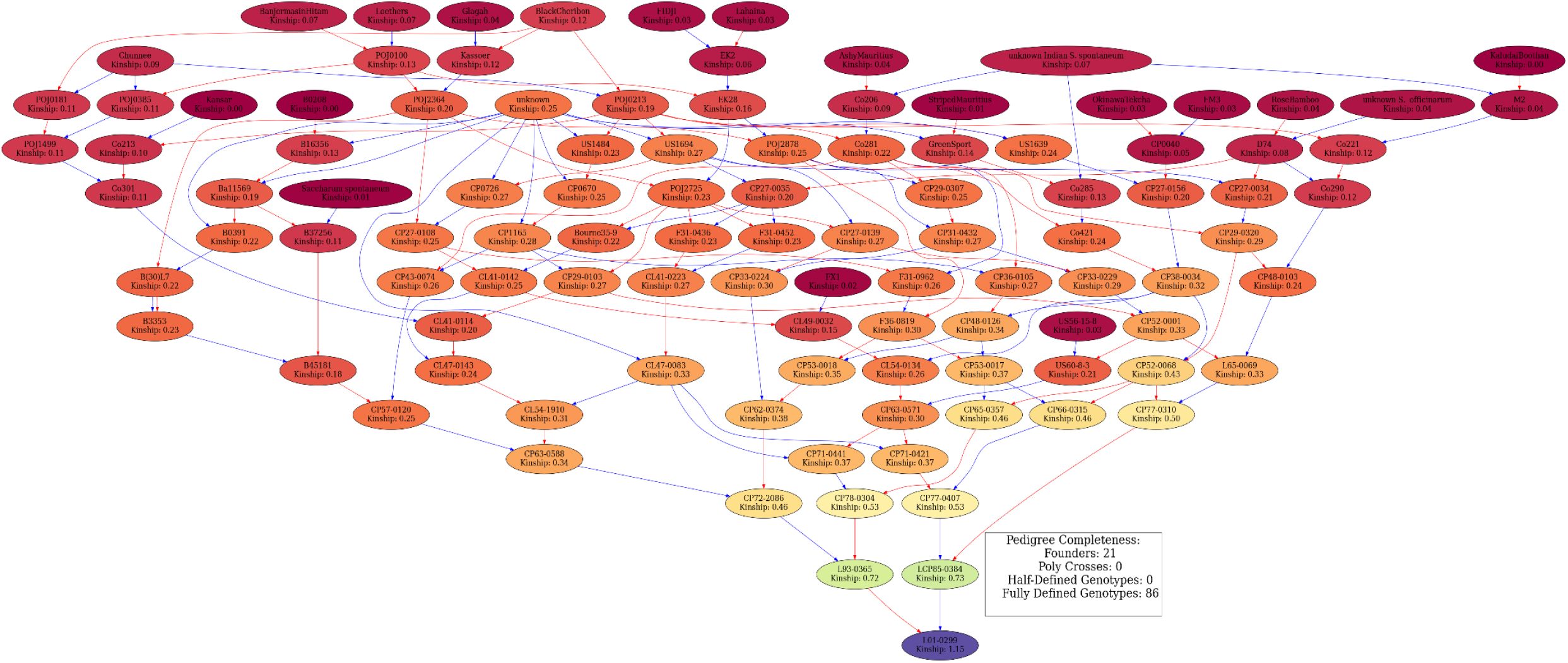
Family tree of L01-0299. Family tree diagrams are generated using graphviz. All pairwise relationships between the individual of interest (L01-0299) and its ancestors are displayed in the individual nodes as well as the name of the genotype. Edges are red for maternal relationships and blue for paternal. A heatmap color scheme is also employed to assist in visualizing the relatedness of the individuals. Pedigree completeness is displayed in the form of a count of founders (individuals with no parents in the dataset), Poly Crosses (for sugarcane genotypes these are instances where the male parent is not known, and these are omitted for data consistency.), Half-Defined genotypes (individuals with only one listed parent), and Fully Defined genotypes (individuals with both parents listed).

Overlapping lineages between two individuals can also be visualized without appending pedigree entries using the Generate Combined Family Trees module. This allows for fast visualization of the shared and unique ancestral lines in the pedigree backgrounds of two individuals. An example of this is shown in **Figure 5**. This functionality enables breeders to have a more visual representation of the relatedness of two individuals, and the ancestors from whom their relatedness originates.

**Figure 5.**
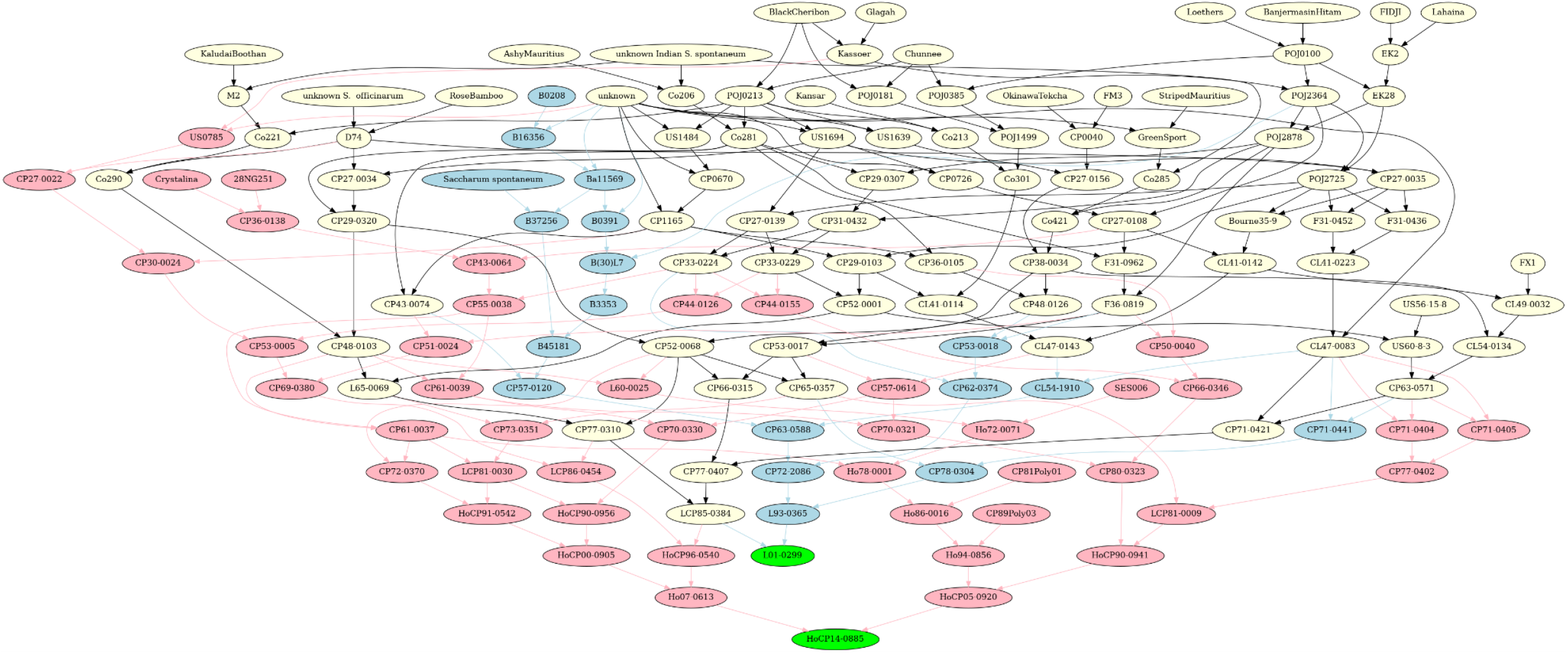
Combined Family Trees of L01-0299 and HoCP14-0885. Pictured is a combined family tree diagram containing two genotypes of interest L01-0299 and HoCP14-0885, highlighted in green. Red nodes and edges are only present in the lineage of HoCP14-0885 and blue nodes and edges are only present in the lineage of L01-0299, yellow nodes and black edges are mutual members of both genotypes lineage.

### Pedigree Search

Apart from generating the family tree and additive kinship matrix, *Canecestry* offers more tools for exploratory analysis of pedigree data. Progeny of individual genotypes can be searched and listed; this also pulls up the second parent of the progeny. Two parental lines can be input to search the dataset for any progeny from that specific crossing event. These functions provide both graphical visualization and a list-based format, putting customization in the hands of the user. A comprehensive list of these features is included in **Figure 6**.

**Figure 6.**
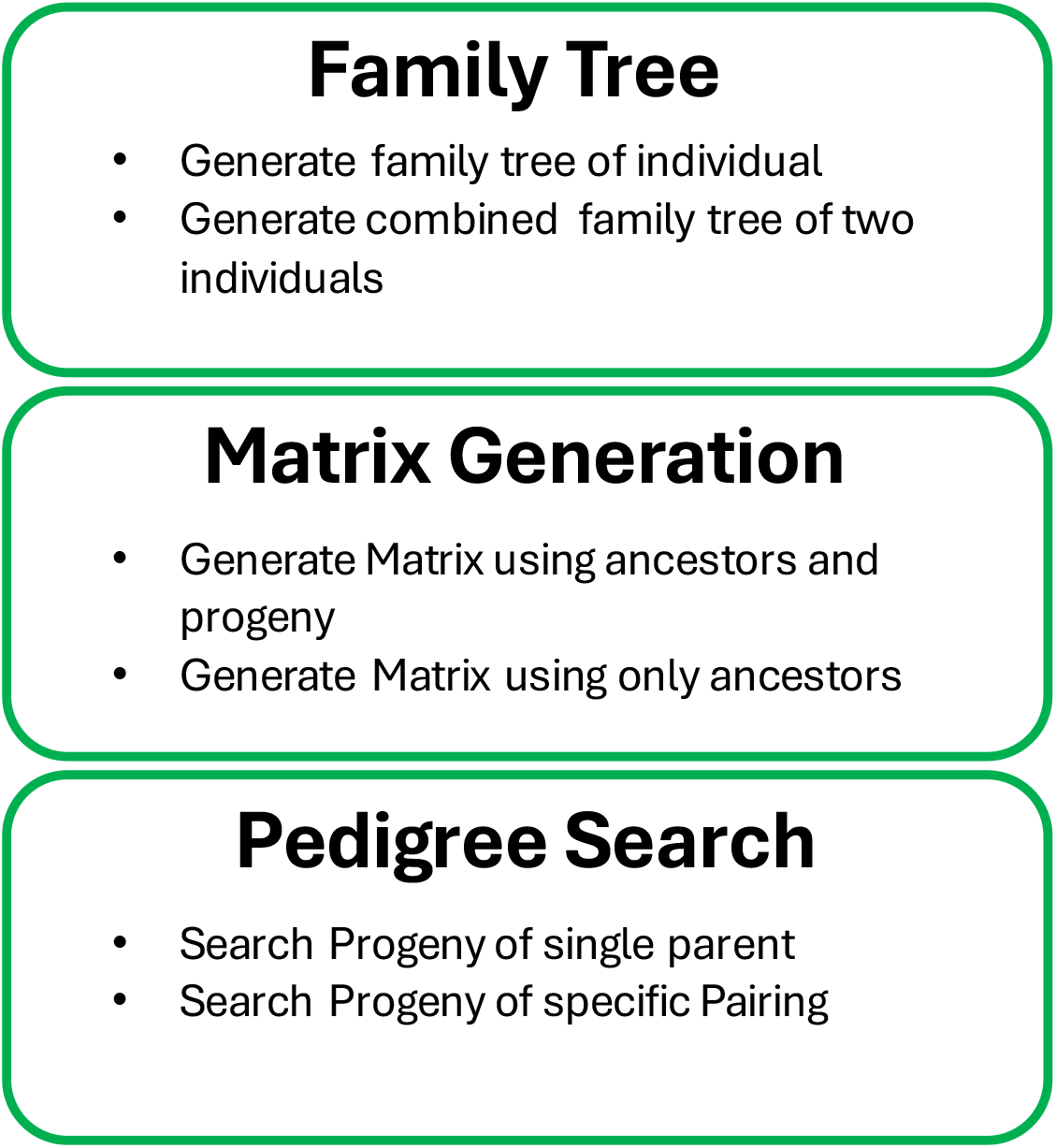
Comprehensive list of tools available for users to visualize subset pedigree information within Canecestry.

### Appending Temporary Pedigree Entries

*Canecestry* employs a dcc.Store method as outlined in the dash web app framework to keep uploaded data persistent across all modules of the application. This allows for data to be appended in app, without having to first modify and reupload data. Pre-existing sugarcane data built in is the primary target for this functionality, as it provides a convenient method for breeders to upload a list of crosses into the dataset to subsequently calculate a pedigree relationship matrix between the progeny. Uploaded entries must be in the same format as the existing dataset, and the male and female parents must have entries as lines in the preexisting dataset. If no overlap is detected, users can manually select the parents from the list of individuals existing in the original dataset. Once added, the updated dataset can be used throughout all modules in *canecestry* until removed using the “trash” button. This button resets the stored dataset back to its previous state.

## Results and Discussion

In this study we introduce Canecestry, a web-based software suite for analysis of complex pedigree datasets. The two primary functions of the application are efficient calculations of additive relationship matrices and the generation of family trees. These functions are broken down into separate modules depending on user need.

Additive numerator relationship matrices can be calculated with the recursive method as outlined by Henderson 1976 or divided by 0.5 to represent the coefficient of coancestry between individuals. Methods are provided to guide the selection of individuals desired by the end user to be included in the calculation to shorten unnecessary computational load. Heatmaps with dendrograms are provided to allow users to visualize the structure of the full matrix that they have calculated. These large matrices can be further subset to only include the relationships that the user needs for subsequent analysis.

Family tree visualizations can be generated for individuals within a user’s input dataset or appended. Kinship values are appended to the nodes to show the relationship between the specified individual and each relative. These graphs can also be generated to highlight distinct ancestral groups between two individuals. Graphs can be subset by generation depth based on user need, with kinship values retaining the full value from the entire lineage. Progeny of a parent or a pairing can be visualized as a graphical list of individuals as well, which can provide an on-hand list of current offspring.

While this application can be utilized by any user with the required data, special consideration has been placed on sugarcane breeders and the intricacies of their pedigree datasets (polycross exceptions). The backbone of this software is a subsetting algorithm that subsets the dataset to only run calculations on necessary individuals to provide fast and convenient insights into pedigree information without the need for data preprocessing, which is lacking in many existing platforms/software.

## Availability and further directions

*Canecestry* is available at canecestry-zanthoxylum2117.pythonanywhere.com. Instructions for installation via Docker or Github are available at https://github.com/Zanthoxylem/CaneCestry

This web application has been designed to be an exploratory tool for pedigree data, with sugarcane breeding as its core user base. Abundant genomic information in other crop breeding programs [28] has allowed for the use of a (G) matrix, or genomic relationship matrix, for genomic-based prediction. These methods take an input of a matrix of individuals and the presence or absence of molecular markers of interest [7]. Methods to calculate these values have been developed for diploid and polyploid species. Pedigree information can often be skewed by biological as well as anthropogenic errors and the use of the (G) matrix to correct these errors has been employed in other crops [29]. The driving force behind *Canecestry* is ease of use, regardless of coding background. Implementation of (G) matrix calculation would provide a user-friendly, web-based interface for calculation, which is currently not present in any packages or software at the time of writing. Both genetic relationship (G) matrices, and pedigree relationship matrices (A) can be combined into a hybrid (H) matrix leveraging the information present in both component matrices. However, computation of this hybrid matrix involves cross validation to determine the most optimal amount of shrinkage to apply when combining the information, placing this type of analysis out of the scope of CaneCestry. Molecular marker datasets can be large, leading to a high computational load. This would require more server-side computing power. Genomic relatedness values would be stored, allowing users to append the (A) matrix and the (G) matrix values onto family tree displays. Displaying this information would give users a better visualization of the relationships between their individuals of interest.

Further applications for *Canecestry* would be its inclusion in an “all in one” software suite to assist breeders in planning and making crosses, incorporating phenotypic trait information as well. Seamless calculation of kinship matrices of potential crosses to see a dynamically updated list of the relatedness within (to aid in parent selection) and between planned crosses within an application for crossing would allow for rapid identification of genetics that are being underutilized. A workflow containing pedigree matrix generation, followed by breeding value estimation would provide rapid insights into daily crossing decisions, with the modular nature of *Canecestry* putting this valuable information in the hands of all breeders regardless of coding knowledge.

## Author Contributions

**Conceptualization: Zachary Taylor, Brayden Blanchard, Ajay Dhungana, Collins Kimbeng Data**

**Curation: Zachary Taylor**

**Funding Acquisition: Collins Kimbeng**

**Software: Zachary Taylor**

**Supervision: Collins Kimbeng**

**Visualization: Zachary Taylor**

**Writing – Original Draft Preparation: Zachary Taylor**

**Writing – Review & Editing: Zachary Taylor, Brayden Blanchard, Ajay Dhungana, Collins Kimbeng**

## Notes

### Competing Interest Statement

The authors have declared no competing interest.

